# TRPV1 in arteries enables a rapid myogenic tone

**DOI:** 10.1101/2021.02.25.432719

**Authors:** Thieu X. Phan, Hoai T. Ton, Hajnalka Gulyás, Róbert Pórszász, Attila Tóth, Rebekah Russo, Matthew W. Kay, Niaz Sahibzada, Gerard P. Ahern

## Abstract

Arterioles maintain blow flow by adjusting their diameter in response to changes in local blood pressure. In this process called the myogenic response, a vascular smooth muscle mechanosensor controls tone predominantly through altering the membrane potential. In general, myogenic responses occur slowly, reaching a plateau in minutes. In the heart and skeletal muscle, however, myogenic tone is rapid; activation occurs in tens of seconds and arterial constrictions or raised extravascular pressure as brief as 100 ms remove tone. Previously, we identified extensive expression of TRPV1 in the smooth muscle of arterioles supplying skeletal muscle, heart and the adipose. Here we reveal a critical role for TRPV1 in the myogenic tone of these tissues. TRPV1 antagonists dilated skeletal muscle arterioles *in vitro* and *in vivo*, increased coronary flow in isolated hearts, and transiently decreased blood pressure. All of these effects of TRPV1 antagonists were abolished by genetic disruption of TRPV1. Stretch of isolated vascular smooth muscle cells, or raised intravascular pressure in arteries (with or without endothelium), triggered Ca^2+^ signaling and vasoconstriction. The majority of these stretch-responses were TRPV1-mediated, with the remaining tone being inhibited by the TRPM4 antagonist, 9-phenantrol. Notably, tone developed more quickly in arteries from wild-type compared with TRPV1-null mice. Furthermore, the rapid vasodilation following brief constriction of arterioles was also dependent on TRPV1, consistent with a rapid deactivation or inactivation of TRPV1. Pharmacologic experiments revealed that membrane stretch activates a phospholipase C/protein kinase C signaling pathway to activate TRPV1, and in turn, L-type Ca^2+^ channels. These results suggest a critical role, for TRPV1 in the dynamic regulation of myogenic tone and blood flow in the heart and skeletal muscle.

## Introduction

Arterioles must deliver blood to tissues within a narrow pressure range to enable adequate perfusion without damaging capillaries. These vessels possess the intrinsic capacity to sense changes in local blood pressure and adjust their caliber to stabilize blood flow. This autoregulatory property, known as “the myogenic response” (Bayliss, 1902), helps to maintain tissue perfusion despite fluctuations in blood pressure. Although an intrinsic property of most arterioles, the kinetics of myogenic tone are especially rapid in skeletal muscle and the heart, tissues that receive phasic blood flow and large increases in their blood supply going from rest to exercise. Fast myogenic tone in coronary and skeletal muscle arterioles may favor uniform spatiotemporal perfusion (Davis, 2012; Goodwill *et al.*, 2017). Further, the rapid release of myogenic tone upon initial skeletal muscle contraction, augments blood flow to improve the transition to work (Clifford, 2007; Davis, 2012). Significantly, impaired or diminished myogenic tone is a pathophysiological feature that accompanies diabetes, obesity (Hodnett & Hester, 2007), coronary artery disease (Duncker *et al.*, 2015), hypertrophic cardiomyopathy (Petersen *et al.*, 2002) and ageing (Lott *et al.*, 2004; Ghosh *et al.*, 2015). All these conditions are associated with declining cardiac vasodilator reserve and exercise performance.

Although Bayliss described the myogenic effect in skeletal muscle more than a century ago (Bayliss, 1902), details of the molecular components are only just emerging. Mechano-activated ion channels, such as Piezo1 are present in vascular smooth muscle cells but do not contribute to normal myogenic tone (Retailleau *et al.*, 2015). Rather, several studies support a role for G-protein coupled receptors (GPCRs) in mechanosensation (Zou *et al.*, 2004; Mederos y Schnitzler *et al.*, 2008) and as vascular smooth muscle mechanosensors (Mederos y Schnitzler *et al.*, 2008; Gonzales *et al.*, 2014; Hong *et al.*, 2016). These GPCRs couple to either a G12/G13 Rho kinase-dependent pathway (Chennupati *et al.*, 2019) or a Gq/G11 dependent phospholipase C (PLC) pathway (Osol *et al.*, 1993; Matsumoto *et al.*, 1995). PLC-mediated signaling activates transduction channels and ultimately voltage-gated Ca^2+^ entry to trigger smooth muscle contraction (Knot & Nelson, 1998; Earley & Brayden, 2015). The identity of the transduction channels is unclear but Transient Receptor Potential (TRP) channels are potential candidates; TRPC6 (Welsh *et al.*, 2002) and TRPM4 (Earley *et al.*, 2004; Reading & Brayden, 2007) channels are implicated in the tone of cerebral arterioles, while TRPP1 is essential for myogenic tone in large-diameter, primary arteries (Bulley *et al.*, 2018). These findings support the existence of distinct, tissue-specific molecular pathways for regulating myogenic tone.

TRPV1 is an ion channel highly expressed in sensory ganglia with important roles in transducing temperature, pain and itch sensory modalities. Heat activates TRPV1, as well as capsaicin, the pungent component of chilli peppers, protons and a variety of endogenous lipid metabolites (Caterina *et al.*, 1997; Caterina & Julius, 2001; Bang *et al.*, 2010). TRPV1 senses and integrates these physical and chemical stimuli via distinct domains that are allosterically coupled to channel gating (Tominaga *et al.*, 1998; Brauchi *et al.*, 2004; Matta & Ahern, 2007). Notably, TRPV1 has been identified in isolated segments of the vasculature (Lizanecz *et al.*, 2006; Kark *et al.*, 2008; Cavanaugh *et al.*, 2011; Czikora *et al.*, 2012; Czikora *et al.*, 2013; Phan *et al.*, 2016). Recently, we performed functional mapping of TRPV1 throughout the mouse circulation. This analysis revealed extensive TRPV1 expression in the vascular smooth muscle of arterioles in the skeletal muscle, heart and adipose tissues (Phan *et al.*, 2020). Furthermore, we found that activation of arteriolar TRPV1 channels by capsaicin or the endogenous lipid, LPA, markedly increased vascular tone and blood pressure (Phan *et al.*, 2020). Of note, the restricted expression of TRPV1 in small arterioles of skeletal muscle matches the classical studies mapping myogenic tone in the limb and skeletal muscle arterial tree (Uchida & Bohr, 1969). Here using *in vitro* and *in vivo* approaches we reveal that TRPV1 mediates the majority of myogenic tone in skeletal muscle and coronary arteries. Further, we reveal that TRPV1 is critical for rapid and dynamic changes in tone in these tissues.

## Methods

### Ethical approval

All procedures were approved by Georgetown University, IACUC Protocol Numbers: 2016-1310 and 2018-0033; George Washington University, **IACUC** number **A202**, and University of Debrecen, Ethics Committee on Animal Research:2/2013/DEMÁB and 4-1/2019/DEMÁB. All efforts were made to minimize the number and suffering of the animals used in this study.

### Animals

Wistar and Sprague-Dawley rats (250-450 g) and C57Bl6 mice (25-30 g) were housed at 24-25°C and had *ad libitum* access to a standard laboratory chow and water.

### Mouse lines

The TRPV1-Cre transgenic mouse line (donated by Dr. Mark Hoon, NIH) was created using a BAC transgene containing the entire TRPV1 gene/promoter (50 kbp of upstream DNA) and IRES-Cre-recombinase (Mishra *et al.*, 2011). Importantly, Cre expression in this mouse faithfully corresponds with the expression of endogenous TRPV1. The TRPV1-Cre (hemizygous) mice were crossed with ai9 ROSA-stop- tdTomato mice (The Jackson Laboratory). The TRPV1^PLAP-nlacZ^ mice (Jackson Laboratory) were developed by Allan Bausbaum and colleagues (UCSF) to express human placental alkaline phosphatase (PLAP) and nuclear lacZ under the control of the TRPV1 promoter (Cavanaugh *et al.*, 2011). The targeting construct contains an IRES-PLAP-IRES-nlacZ cassette immediately 3’ of the TRPV1 stop codon, which permits the transcription and translation of PLAP and nlacZ in cells expressing TRPV1 without disrupting the TRPV1 coding region. TRPV1-null mice were purchased from The Jackson Laboratory. TRPV1-Cre:ChR2/tdTomato mice were generated by crossing TRPV1-Cre mice with ChR2/tdTomato mice (The Jackson Laboratory).

### mRNA analysis

Mice were anesthetized with isoflurane (4% in 100% O2) and euthanized by perfusing the heart with ice-cold PBS (0.1 M, pH 7.3). Dorsal root ganglia (L4, L5), brachial and radial branch arteries were isolated. RNA was extracted and purified using the RNAqueous Micro Kit (Invitrogen). First-strand cDNA synthesis was performed using SuperScript III Reverse Transcriptase (Invitrogen) with the supplied oligo(dT)20 primer. The resulting cDNA was used as a template for PCR amplification.

### Arterial smooth muscle (ASM) cell isolation

Adult mice (C57BL/6J) were euthanised by CO_2_ /decapitation. Radial branch and cerebellar branch arteries were washed in Mg^2+^-based physiological saline solution (Mg-PSS) containing 5 mM KCl, 140 mM NaCl, 2 mM MgCl_2_, 10 mM Hepes, and 10 mM glucose (pH 7.3). Arteries were initially digested in papain (0.6 mg/ml) (Worthington) and dithioerythritol (1 mg/ml) in Mg-PSS at 37°C for 15 min, followed by a 15-min incubation at 37°C in type II collagenase (1.0 mg/ml) (Worthington) in Mg-PSS. The digested arteries were washed three times in ice-cold Mg-PSS solution and incubated on ice for 30 min. After this incubation period, vessels were triturated to liberate smooth muscle cells and stored in ice-cold Mg-PSS before use. Smooth muscle cells adhered loosely to glass coverslips and were studied within 6 hours of isolation.

### Ca^2^ imaging

ASM cells and arteries were respectively loaded with 5 μM and 10 μM Fluo-4-AM (Invitrogen, Thermo Fisher Scientific) in a buffer solution containing 140 mM NaCl, 4 mM KCl, 1 mM MgCl2, 1.2 mM CaCl2, 10 mM HEPES, and 5 mM glucose (pH 7.3). Temperature was maintained at 32-33°C using a heated microscope stage (Tokai Hit). Bath temperature was verified by a thermistor probe (Warner instruments). ASM cells and arteries were imaged with 10X and 20X objectives using a Nikon TE2000 microscope with an excitation filter of 480 ± 15 nm and an emission filter of 535 ± 25 nm. The images were captured by a Retiga 3000 digital camera (QImaging) and analysis was performed offline using ImageJ.

### Ex vivo artery physiology

Mice and rats were euthanized (CO_2_/decapitation). Skeletal muscle arteries (radial artery branch, artery #18, subscapular branch artery #14, see Fig. 6G, (Phan *et al.*, 2020)) were isolated and cannulated with glass micropipettes, and secured with monofilament threads. In some experiments arteries were denuded of endothelium by passing 1 ml of air followed by 1 ml of PSS through the lumen. Effective removal of the endothelium was confirmed by the absence of dilation of the arteries to ACh. The pipette and bathing PSS solution containing (in mM): 125 NaCl, 3 KCl, 26 NaHCO_3_, 1.25 NaH_2_PO_4_, 1 MgCl_2_, 4 D-glucose, and 2 CaCl_2,_) was aerated with a gas mixture consisting of 95% O_2_, 5% CO_2_ to maintain pH (pH 7.4). To induce maximal dilation, arteries were perfused with a PSS solution containing 0 CaCl_2,_ 0.4 mM EGTA and 100 *μ*M sodium nitroprusside (SNP). Arterioles were mounted in a single vessel chamber (Living Systems Instrumentation) and placed on a heated imaging stage (Tokai Hit) to maintain bath temperature between 34-35°C, while intraluminal pressure was maintained by a Pressure Control Station (Stratagene) at 60 mmHg. Arteries were viewed with a 10X objective using a Nikon TE2000 microscope and recorded by a digital camera (Retiga 3000, QImaging). The arteriole diameter was measured at several locations along each arteriole using the NIH-ImageJ software’s edge-detection plug-in (Diameter) (Fischer *et al.*, 2010). The software automatically detects the distance between edges (by resampling a total of five times from adjacent pixels) yielding a continuous read-out ±SD of a vessel’s diameter.

### Intravital imaging

Intravital imaging was performed in radial artery branches (about 60 *μ*m in diameter, artery #18 in Fig 6G, (Phan *et al.*, 2020)). Mice were restrained by grasping the skin at the nape of the neck and anesthetized with urethane (1.2 g/kg/IP). The adequacy of anesthesia was confirmed by the absence of pedal and corneal reflexes. The forelimb was shaved and an incision was made. The skin and underlying muscle tissue were reflected to expose the brachial-radial artery junction. Both in WT and TRPV1-null mice, the arteries were visualized with a Zeiss stereomicroscope and illuminated with a low power blue light (using a standard GFP filter cube) exploiting the differential auto-fluorescence between tissue and blood. In TRPV1-Cre:ChR2/tdTomato mice, arteries were visualized with low power visible irradiation and stimulated with blue light. The exposed arteries were locally perfused (using a 250 *μ*m cannula connected to a valve-controlled gravity-fed perfusion system) with preheated buffer described for Ca^2+^ imaging. The surface tissue temperature (34-35°C) was measured via a thermistor (Warner Instruments) that was positioned next to the artery. Arteries were challenged with buffer without Ca^2+^ and with 1 mM EGTA to measure the passive diameter. The arteriole diameter was measured using ImageJ as described above for the *ex vivo* vessels. After the recordings, the mice were euthanized (CO_2_/decapitation).

### Coronary flow measurements

Sprague–Dawley rats (male, 300–350 g) were placed in a deep surgical plane of anesthesia by isoflurane inhalation (4% in 100% O_2_), confirmed by lack of pedal reflex. The heart was then exposed via thoracotomy, quickly excised and rinsed in a bath of ice-cold perfusate. The aorta was rapidly cannulated then flushed with 500 units of heparin mixed with the perfusate that contained (in mM): 118 NaCl, 4.7 KCl, 1.25 CaCl_2_, 0.57 MgSO_4_, 1.17 KH_2_PO_4_, 25 NaHCO_3_, and 6 glucose. Hearts were then transferred to a retrograde perfusion system that delivered oxygenated (gassed with 95%O_2_-5%CO_2_) perfusate to the aorta at constant pressure (70mmHg) and 37±1°C. Coronary flow was measured using a tubing flowsensor (Transonic Systems) placed above the aortic cannula and was continuously acquired with the ECG using a PowerLab system (ADInstruments). BCTC was added to the perfusate reservoir. Data were analyzed off–line and the integral of coronary flow was calculated between the time of injection and the onset of the hyperemia response.

### Sensory nerve ablation

Neonate TRPV1^PLAP-nlacZ^ mice were anesthetized briefly (5 min) with isoflurane (4-5% in 100% O_2_) and treated with resiniferatoxin (50 ug/kg s.c.) at postnatal days 2 and 5. Animals were allowed to recover in a warm environment (30°C, 2 hr) to minimize any hypothermic effects. This dose of resiniferatoxin causes profound sensory nerve desensitization/block and mice exhibited no outward signs of distress in recovery, and there was no disruption to dam-pup interactions. At 8-12 weeks, mice were either euthanized for tissue collection, or used for BP measurements followed immediately by euthanasia (CO_2_/decapitation). Sensory nerve ablation was confirmed by nuclear LacZ staining of DRG ganglia.

### Systemic blood pressure recording

The experiments were performed in anesthetized mice (urethane 1.2-1.5 g/kg/IP) and rats (thiopental 50 mg/kg/IP; supplemented by 5 mg/kg/IV if needed). After anesthesia, mice or rats underwent cannulation of the carotid artery and jugular vein as follows:

#### Surgical preparation in the mouse

After the depth of anesthesia was confirmed by lack of pedal and corneal reflexes, mice were intubated via the trachea after tracheotomy to maintain an open airway and to institute artificial respiration when necessary. Next, the left carotid artery and the right jugular vein were cannulated with a Millar catheter (1F SPR-1000) and a polyethylene tubing (PE-10), respectively, for monitoring arterial blood pressure and for systemic (intravenous) infusion of drugs. To monitor heart rate, a three-point needle electrode-assembly representing Lead II of the electrocardiogram (ECG) was attached subcutaneously to the right and left forelimbs along with a reference electrode to the left hindlimb. Both the Millar catheter and the ECG assembly were coupled to a PowerLab data acquisition system (ADInstruments). Before vessel cannulation, the adjacent left cervical vagus was carefully isolated from the left carotid artery. Body temperature was monitored by a digital rectal thermometer and maintained at 37 ± 1°C with an infrared heat lamp. After the study, mice were euthanized (CO_2_/decapitation).

#### Conscious blood pressure recordings

RTX-treated mice (8-12 weeks) were anesthetized with isoflurane (2-4%) and surgically implanted with in-dwelling jugular catheters (Instech labs., USA). The adequacy of anesthesia was confirmed by the absence of pedal and corneal reflexes. Post-surgically mice were administered carprofen (one dose of 5mg/kg s.c.) and placed in a cotton-filled cage until ambulatory. After 72 h, the animals were prepared for BP recording. BP was measured by tail-cuff plethysmography (Coda6, Kent Scientific, USA) performed before and immediately after the infusion of drugs. At the end of the study, the mice were euthanized (CO_2_/decapitation).

#### Surgical preparation in the rat

Before the commencement of surgical interventions, the depth of anaesthesia was checked by squeezing the tip of the rat’s tail. If no response to this challenge was observed, the animal was fixed to a plastic foil-covered polystyrene plate by strings and tapes. The animal was placed in a supine position and the collar region was shaved by a razor. A midline incision was made to expose the trachea, carotid arteries, and the jugular vein. Similar to the mouse, following intubation of the trachea, the left carotid artery, and jugular vein were cannulated with a polyethylene tubing (PE50) to monitor blood pressure and infuse drugs, respectively. Blood pressure (and ECG) was continuously recorded via a pressure transducer connected to the Haemosys hemodynamic system (Experimetria, Budapest, Hungary). The ECG was recorded from the extremities of the animal using hypodermic metal needles inserted subcutaneously per the Einthoven method (I, II, III leads). As in the mouse, heart rate was determined from lead II of the ECG recordings, and body core temperature was maintained at 37±1 °C with a temperature-controlled infrared heating lamp. During the experiment, the depth of anesthesia was checked regularly and if necessary was supplemented by an intravenous dose of 5 mg/kg Thiopental. After the study, the animals were euthanized (CO_2_/decapitation).

#### Drug administration

Intravenous infusion of drugs was initiated only when a stable baseline of blood pressure and heart rate was present. This was also the case when drugs were re-administered. Final drug solutions contained: capsaicin (saline with 0.4% EtOH), LPA (saline with 0.6% EtOH).

#### Chemicals

Capsaicin, resiniferatoxin and 4-(3-Chloro-2-pyridinyl)-*N*-[4-(1,1-dimethylethyl)phenyl]-1-piperazinecarboxamide (BCTC) were purchased from Tocris Bioscience or Adooq Bioscience and stock solutions were prepared in EtOH at 1 M and 100 mM, respectively. Lysophosphatidic acid (LPA) C18:1 was purchased from Cayman Chemical. Unless otherwise indicated, all other chemicals were obtained from Sigma–Aldrich.

#### Statistical analysis

Data were analyzed using Prism (GraphPad Software, La Jolla, CA) and are expressed as means ± SD. Unless otherwise stated, statistical significance was evaluated using t-test and one-way ANOVA with treatment interactions assessed by Tukey’s *post hoc* multiple comparisons test. A *P* value of < 0.05 was considered statistically significant.

## Results

### TRPV1 contributes to resting tone in skeletal muscle and coronary arterioles

Previously, we demonstrated prominent TRPV1 expression in arterioles supplying skeletal muscle, heart and adipose tissues (Phan *et al.*, 2020). To identify potential physiological roles for TRPV1 in regulating vascular tone we tested the effects of the highly specific TRPV1 antagonists, BCTC and AMG517, in pressurized arterioles isolated from mouse skeletal muscle. Notably, bath application of BCTC (1 μM) or AMG517 (10 μM) rapidly dilated wild-type arterioles by ~35% (P = 0.0001 and P= 0.0016, Fig. 1A-D). Importantly and in contrast, the antagonists did not affect the diameter of arteries obtained from TRPV1-null mice (P>0.9999, Fig. 1A-D) indicating that their dilatory effects were mediated by on-target pharmacologic inhibition of TRPV1. Next, to test for a contribution of TRPV1 to *in vivo* vascular tone, we performed intravital imaging of skeletal muscle feed arterioles (~60 *μ*M diameter) supplying the forelimb radial muscles (Fig. 2). Here, we locally perfused BCTC while maintaining the tissue temperature at 34±0.5°C. Similar to the results in isolated arterioles, BCTC dilated *in vivo* arterioles in WT but not in TRPV1-null mice (Fig. 2A and B). The dilation to BCTC exhibited a concentration-dependent relationship with a peak dilatory effect ~70% of that produced by a zero Ca^2+^ solution (Fig. 2C and D). Further, BCTC had a greater dilatory effect in small diameter (~25 μm) compared with medium diameter (~60 μm) arterioles (*P =* 0.002, Fig. 2E), in agreement with larger resting tone in smaller arterioles (Fig 2F).

**Figure 1.**
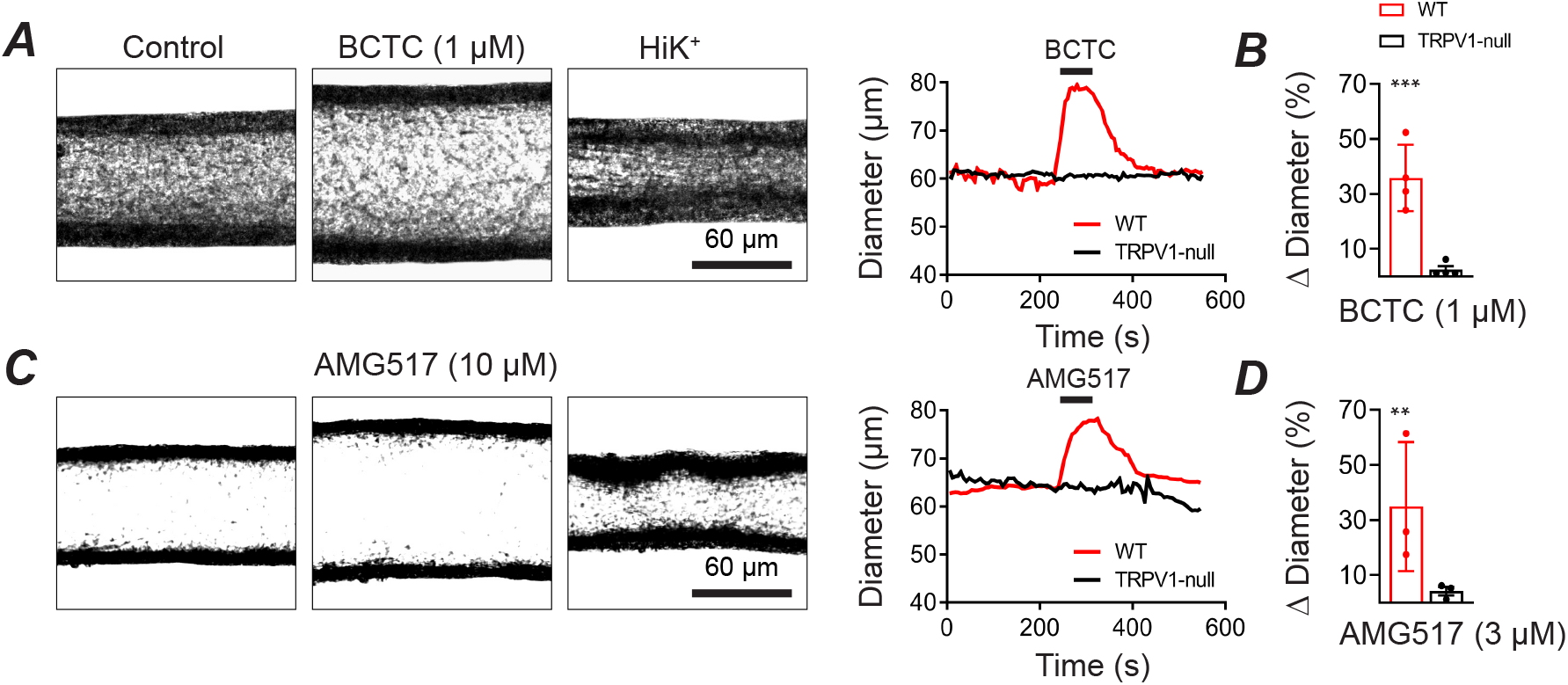
TRPV1 antagonists dilate isolated skeletal muscle arterioles. *A* and *B*, BCTC (1 μM) dilates isolated, pressurized (60 mmHg) skeletal muscle arteries from wild-type (n = 5) but not TRPV1-null (n = 4) mice (*\*\*\*P =* 0.0016). ***C*** and ***D*** AMG517 (3 *μ*M) selectively dilates arteries from WT (n = 3) but not TRPV1-null (n = 3) mice (unpaired t-test, *\*\*\*P =* 0.000114).

**Figure 2.**
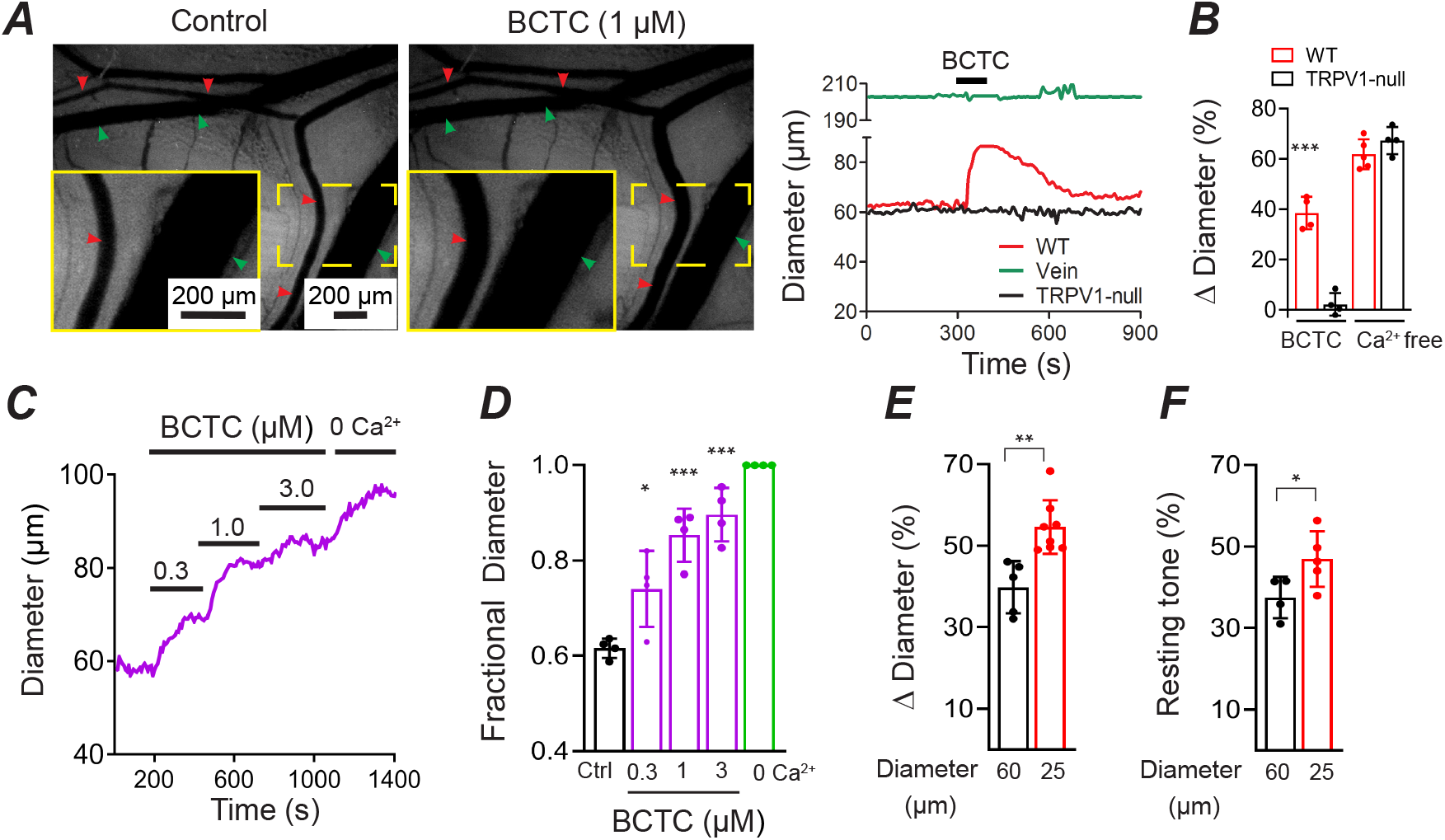
TRPV1 antagonists dilate skeletal muscle arterioles *in vivo*. ***A*,** Intravital imaging show that BCTC (1 μM) dilates *in vivo* radial branch arterioles (red arrows) without affecting arterioles in TRPV1-null mice or veins (green arrows). ***B***, Mean arteriole dilation evoked by BCTC in WT (n = 5) and TRPV1-null mice (n = 5) and maximal dilation with zero Ca^2+^/EGTA (unpaired t-test, \*\*\**P =* 0.000088. ***C*** and ***D***, Concentration-dependent dilation by BCTC normalized to zero Ca^2+^/EGTA (n = 4 arteries, \**P =* 0.024, \*\**P =* 0.00023, \*\*\**P* < 0.0001). ***E*** and ***F***, BCTC-evoked dilation and resting tone in small (~25 μm, n = 8) and medium (~60 μm, n = 5) diameter arteries (unpaired t-test, \**P =* 0.048, \*\**P =* 0.0022).

Next, we tested for the presence of TRPV1-dependent myogenic tone in the heart. Previously, we identified prominent TRPV1 expression in small-diameter, sub-epicardial arterioles (<120 *μ*m diameter) of the ventricular myocardium, and we therefore asked whether TRPV1 contributes to tone in the coronary circulation. Indeed, inline infusion of BCTC into the isolated rat hearts (3 μM final concentration) markedly increased coronary flow by ~600 ml/min.s (*P* = 0.0018, Fig. 3A and B), that partially recovered over a 15-minute period. The elevated coronary perfusion occurred without alterations in heart rate (Fig. 3C). Thus, we conclude that TRPV1 contributes significantly to coronary myogenic tone.

**Figure 3.**
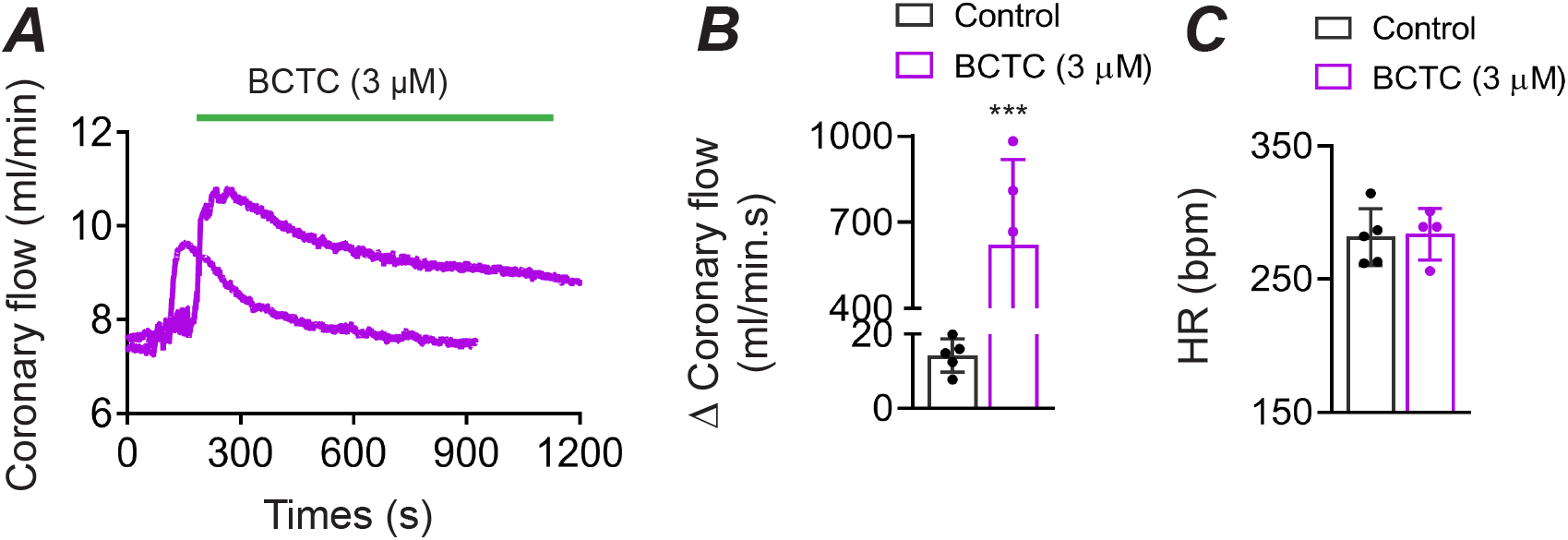
BCTC increases coronary perfusion. ***A***, BCTC infusion into the coronary circulation increases coronary perfusion in an isolated rat heart. ***B***, Mean changes in perfusion evoked by BCTC (measured over 13 to 17 minutes, n = 5, unpaired t-test, *** *P* = 0.0018), ***C***, BCTC does not affect the heart rate (n = 5, unpaired t-test, *P* = 0.874842).

### TRPV1 antagonists transiently decrease systemic blood pressure

Since skeletal muscle arterioles contribute significantly to systemic BP we asked whether inhibition of TRPV1-dependent tone would alter BP. In rats, bolus IV administration of BCTC dose-dependently decreased BP by up to 25 mmHg (Fig. 4A and B). AMG517 also significantly decreased BP (Fig. 4C). In mice, an IV infusion (10 s duration) of BCTC similarly decreased BP by ~15 mm Hg, comparable in magnitude to sodium nitroprusside (50μg/kg, Fig. 4D and E). In contrast, BCTC did not change BP in TRPV1-null mice. Furthermore, in conscious, sensory-nerve ablated mice, we observed equivalent BP responses to BCTC (~15 mmHg, Fig. 4F), ruling out side effects of anesthesia and revealing an arterial delimited effect. The peak responses to BCTC occurred without any significant changes in heart rate (Fig. 4G), however during prolonged (5 min) infusion of BCTC the BP recovered to baseline accompanied by an increase in heart rate (P = 0.0176, Fig. 4H and I).

**Figure 4.**
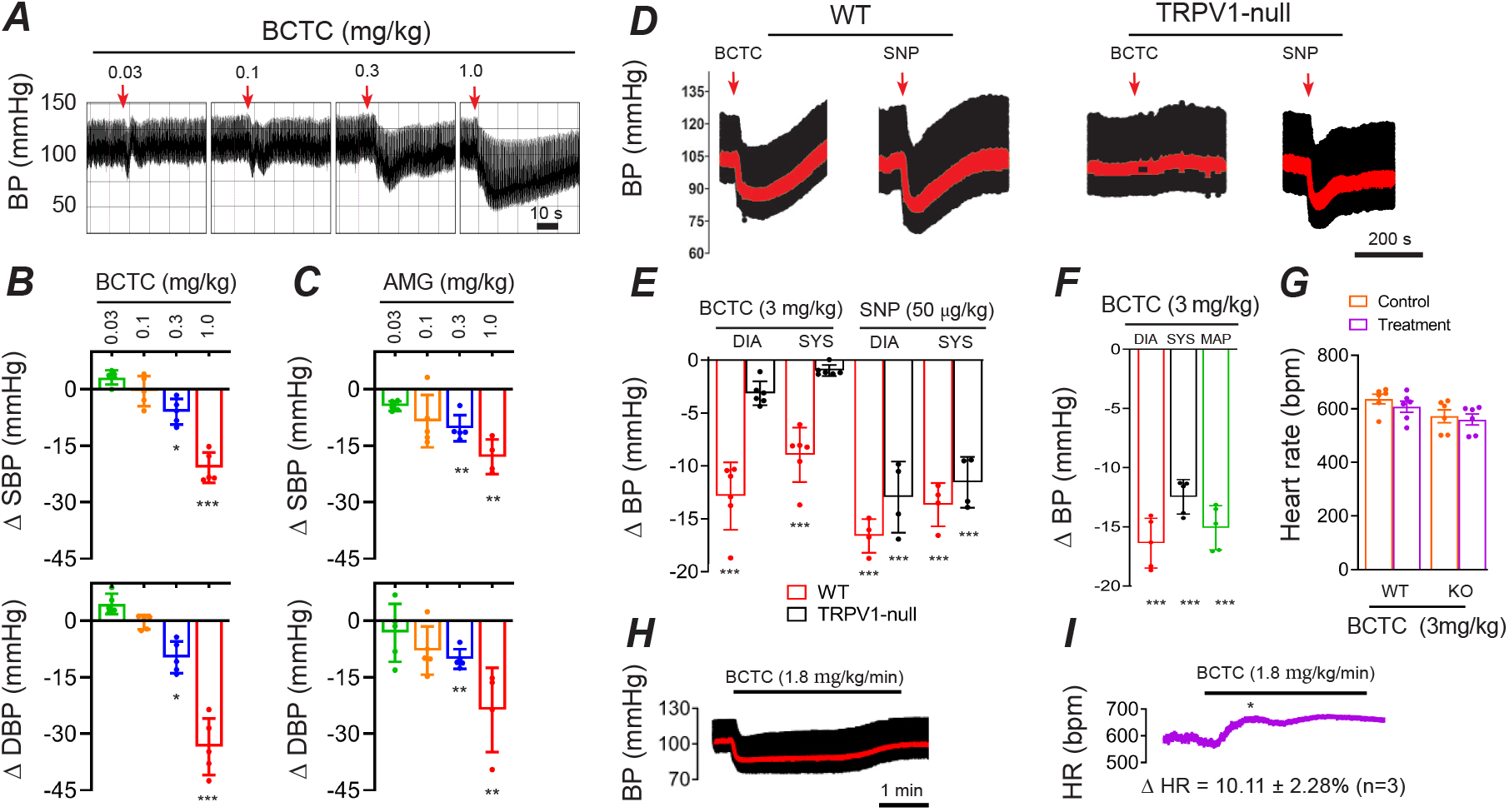
BCTC transiently decreases systemic blood pressure. ***A-C***, Blood pressure (BP) changes in rats during bolus IV BCTC or AMG517 (n = 5, one-way ANOVA, \**P* < 0.05, ** *P* < 0.01, \*\*\**P* < 0.001). ***D*** and ***E***, BP changes in WT and TRPV1-null mice in response to IV infusion (20 s) of BCTC (n = 6) and sodium nitroprusside (n = 4, unpaired t-test, ** *P* < 0.01, \*\*\**P* < 0.001). Mean arterial pressure is shown in red. ***F***, BP changes in response to IV BCTC in conscious, sensory nerve-ablated mice (n = 5, unpaired t-test, *\*\*\*P < 0.001*). ***G*** and ***H***, Representative changes in BP and heart-rate in response to prolonged (4 minute) infusion of BCTC.

### TRPV1 enables a rapid myogenic tone

The dilation of skeletal muscle and coronary arterioles by TRPV1 antagonists suggests a role for TRPV1 in the regulation of myogenic tone in these tissues. To explore this hypothesis directly, we measured the responses of isolated skeletal muscle feed arterioles (~120μm) to changing intraluminal pressure (20 - 100 mmHg). Step increases in pressure immediately increased the arterial diameter followed by reflex constrictions (reflecting developing myogenic tone) that became progressively faster at higher pressure levels (Fig. 5A, D, E). Notably, BCTC markedly inhibited these myogenic constrictions by greater than 50% (P < 0.01). Moreover, while arteries from TRPV1-null mice exhibited an unchanged final tone (recorded after 5 min), at pressures greater than 60 mm Hg the development of tone was significantly slower compared with WT arteries (P < 0.05, Fig. 5B-E). The difference between pharmacologic and genetic disruption of TRPV1 may reflect developmental compensation in the TRPV1-null mice.

**Figure 5.**
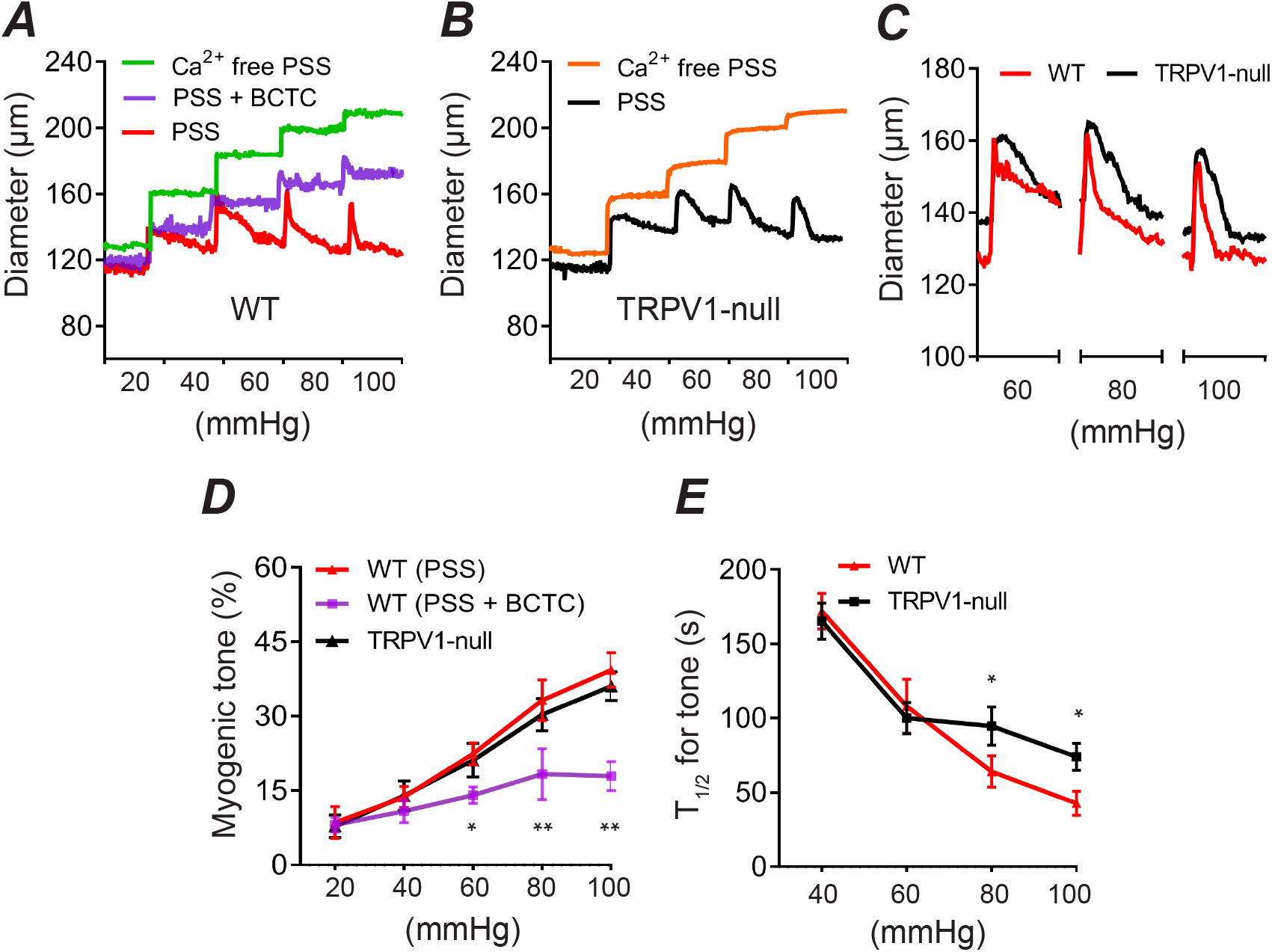
TRPV1 mediates rapid myogenic tone in vitro. ***A-C***, Pressure-diameter relationship in isolated skeletal muscle arterioles from WT and TRPV1-null mice, control (red), 3 μM BCTC (purple) or 0 Ca^2+^/EGTA/SNP (green). Bath temperature was 35°C. ***D*** and ***E***, Myogenic tone and kinetics for development of tone (T_1/2_) versus intraluminal pressure WT (n = 3), WT + BCTC (n = 4) and TRPV1-null (n = 3, one-way ANOVA, \**P* < 0.05, \*\**P* < 0.01).

**Figure 6.**
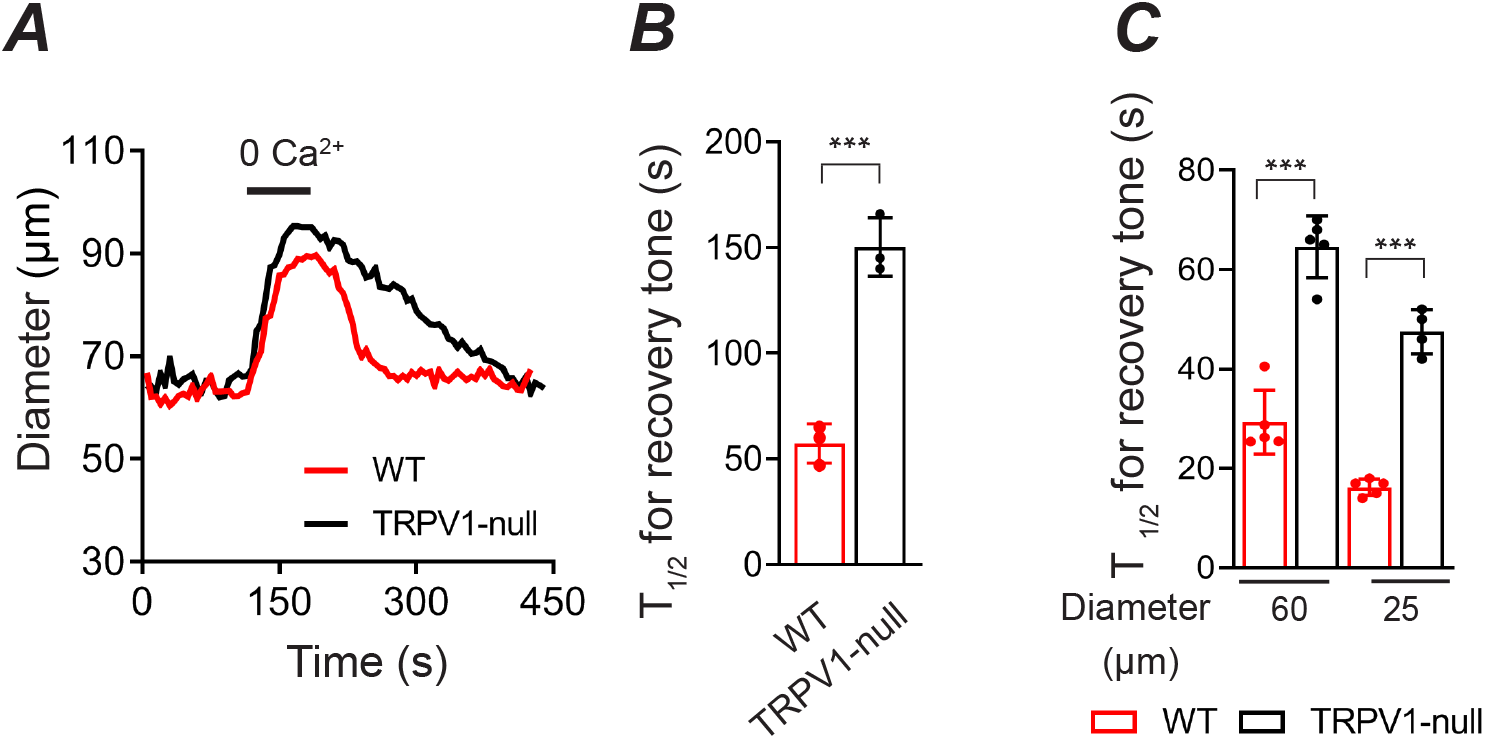
TRPV1 mediates rapid myogenic tone *in vivo*. ***A** and **B***, Development of tone in radial muscle branch arteries after a maximal dilation (0 Ca^2+^/EGTA) in WT and TRPV1-null mice (n = 3, unpaired t-test, \*\*\**P* = 0.0006). ***C***, The time course for recovery of tone after a reactive vasodilation response to KCl in small (n = 5) and medium (n = 5) diameter arterioles form WT and TRPV1-null mice (unpaired t-test, *\*\*\*P < 0.0001*).

To further examine the kinetics of myogenic tone in small (~25 μm) and medium (~60 μm)-diameter arterioles that highly express TRPV1, we turned to intravital imaging. After medium-diameter arterioles *in vivo* were dilated with a zero Ca^2+^ buffer, the tone returned 3-times faster (P = 0.0006) in wild-type compared with TRPV1-null mice (Fig. 6A and B). Similarly, the recovery of tone after a brief vasodilation was 2-3-fold faster in wild-type mice; Figure 6C show that genetic disruption of TRPV1 increased the T_1/2_ for tone from 16 s to 48 s in small diameter arterioles (P = 1.5 ×10^6^) and from 29 s to 65 s in medium diameter arterioles (P = 2×10^5^). Thus, pharmacological or genetic disruption of TRPV1 impairs the magnitude and/or rate of development of myogenic tone.

### TRPM4 contributes to myogenic tone of skeletal muscle arterioles

Previous studies have showed that TRP channels regulate myogenic tone via depolarization-induced activation of the L-type Ca^2+^ channel, Ca_V_1.2 (Knot & Nelson, 1998; Earley & Brayden, 2015). Indeed, we found that the Ca_V_1.2 antagonist, nifedipine, inhibited ~75-80% of tone in skeletal muscle arterioles (Fig. 7E). Further, nifedipine treatment prevented the BCTC-evoked dilation consistent with TRPV1 signaling entirely via the Ca_V_1.2 pathway. Analysis of the respective relaxation to nifedipine and BCTC revealed that TRPV1 signaling contributes approximately two-thirds of the Ca_V_1.2 dependent tone (Fig. 7E). To identify the TRPV1-independent component of Ca_V_1.2-mediated tone we screened WT and TRPV1-null arteries for mRNA expression of candidate channels with known expression in arteries and/or thermosensitivity; TRPC3 & 6, TRPM2, 3 & 4, TRPP1 and anoctamin1. This analysis (Fig.7A) showed that genetic disruption of TRPV1 led to a significant increase (P < 0.05) in TRPM4 and TRPP1 expression. Interestingly, numerous studies have demonstrated a role for TRPM4 in the myogenic tone of different vascular beds, while a recent study showed that TRPP1 contributes to the modest tone found in primary arteries. To test a role for TRPM4 in skeletal muscle we employed the antagonist 9-phenanthrol. Treatment with 9-phenanthrol dilated arteries and reduced tone by 39% (Fig. 7B - D). Further, co-treatment with 9-phenanthrol and BCTC inhibited tone to the same extent as nifedipine (Fig. 7). Notably, 9-phenanthrol produced a significantly greater dilation in arteries from TRPV1-KO compared with WT mice (Fig, 7B-D), an effect consistent with the increased expression of TRPM4 following genetic disruption of TRPV1. This compensatory change, however, did not restore the kinetic properties, and TRPV1 appears critical for the fast development of tone.

**Figure 7.**
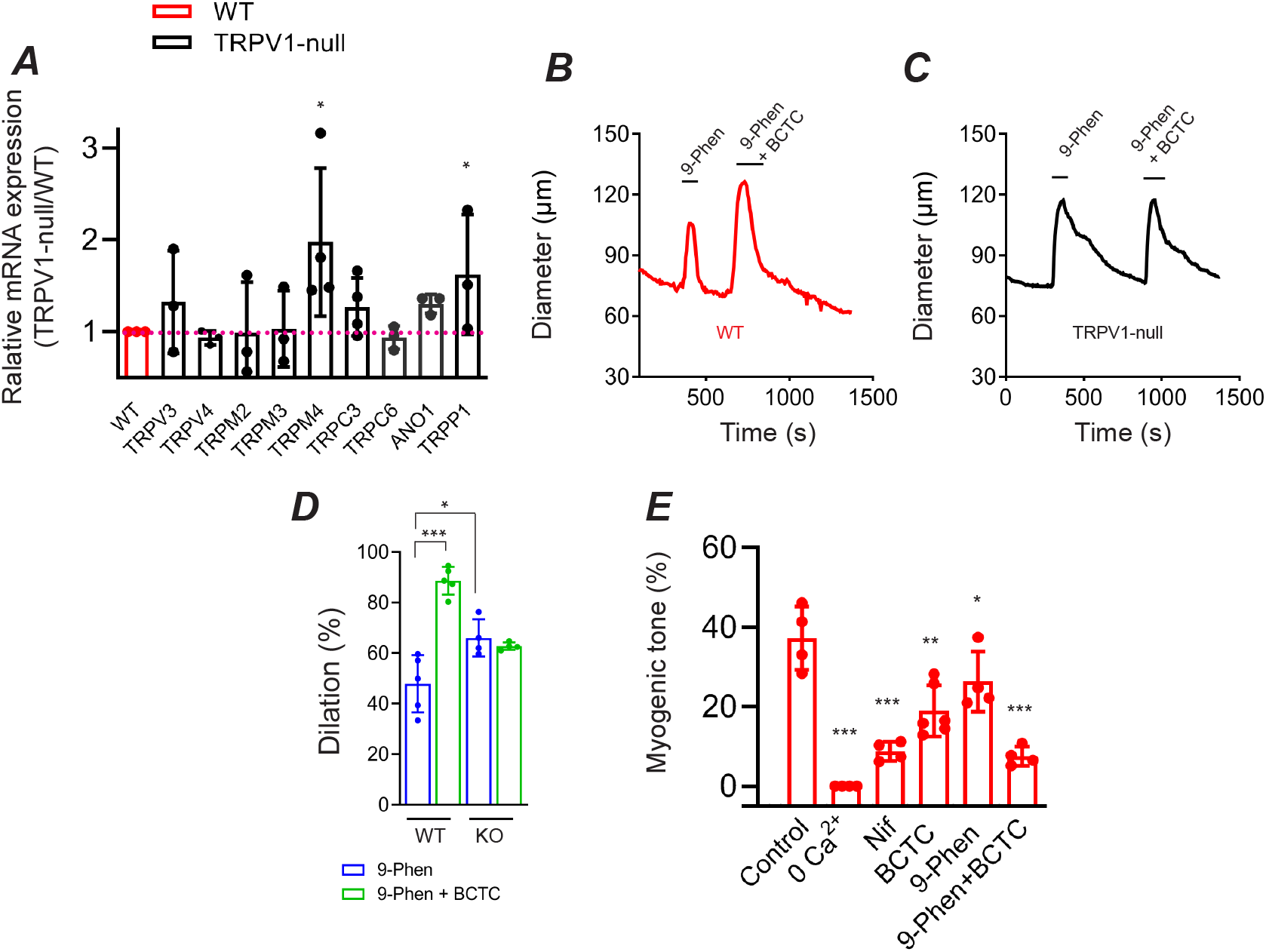
TRPM4 contributes to non TRPV1-dependent myogenic tone. ***A***, Relative mRNA expression for candidate ion channels in TRPV1-null compared with WT skeletal muscle arteries (n = 3 mice, \**P* < 0.05). ***B -D***, Arterial dilation evoked by 9-phenanthrol (30 μM) and 9-phenanthrol plus BCTC in WT (n = 5) and TRPV1-null (n = 4) arteries \*\*\**P* < 0.001, \**P* =< 0.001). ***E*,** Summary of myogenic tone in WT arteries; control (n = 4), 0 Ca2+ (n = 4), nifedipine (5 μM, n = 4), BCTC (3 μM, n = 6), 9-phenanthrol (30 μM, n = 4), BCTC plus 9-phenanthrol (n = 4). \*\*\**P* < 0.001, \*\**P* =< 0.001, \**P=* < 0.001).

### TRPV1 enables rapid reactive vasodilation

At the onset of physical activity, skeletal muscle arterioles immediately dilate (Clifford, 2007), and this phenomenon may involve a relaxation of myogenic tone (Davis, 2012). We therefore tested whether TRPV1 contributes to this process by measuring the dynamic changes in the diameter of skeletal muscle arterioles *in vivo* in response to vasoconstrictive stimuli. We found that local application of KCl (40 s) in wild-type mice or blue light (40 s) in TRPV1-Cre:ChR2 mice constricted arteries followed by a rebound dilation of approximately 35% (Fig. 8A and C). Notably, BCTC fully inhibited the post-KCl dilation (Fig. 8B and C). Furthermore, disruption of *Trpv1* gene expression markedly suppressed (by >70%) the amplitude of the dilation (Fig. 8A and C; *P* < 0.001). Hyperemia following a 40 s arterial constriction may reflect both myogenic and metabolic pathways. However, we observed similar arterial dilations after brief (4 - 6 s) constrictions evoked by either KCl or blue-light pulses (Fig. 8D-F), supporting a primary role for a myogenic mechanism (Davis, 2012). Additionally, both the magnitude of the vasodilation was approximately two-fold greater in small diameter (~25 μm) compared to medium diameter (~60μm) arterioles (Fig. 8D-F). These data suggest that upon arterial constriction TRPV1 is rapidly deactivated or inhibited to promote a rebound vasodilation. Subsequently, after a brief period of hyper-perfusion, TRPV1 reactivates to restore myogenic tone.

**Figure 8.**
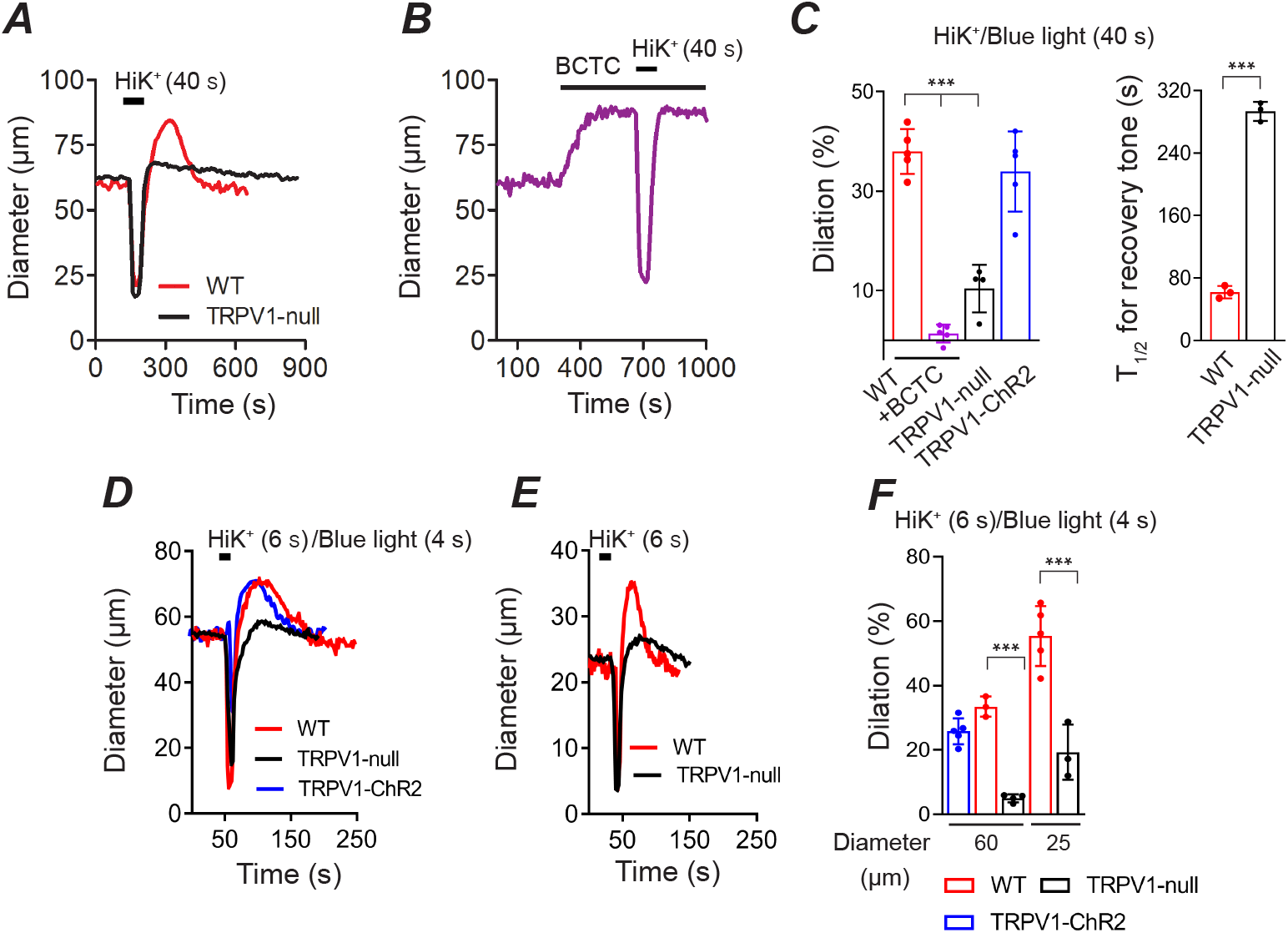
TRPV1 is critical for reactive vasodilation. ***A-C***, Arterial dilation following 40 s constriction evoked by KCl or blue light (TRPV1-ChR2 mice) and the reemergence of tone. BCTC and disruption of the TRPV1 gene inhibits the magnitude of the dilation and rate of recovery of tone (WT, n = 5; WT+BCTC, n = 5; TRPV1-null, n = 4; TRPV1-ChR2, n = 5, \*\*\**P* < 0.001). ***D-F***, Dilation of medium-diameter (~60 μm) and small (~25 μm) arteries in response to 4 s blue light (TRPV1-ChR2 mice) or 6 s application of KCl (WT, n = 3-5; TRPV1-null, n = 3-4; TRPV1-ChR2, n = 5, \*\*\**P* < 0.001).

### A PLC/PKC pathway underlies stretch-evoked activation of TRPV1 in vascular smooth muscle

To explore whether stretch-mediated activation of TRPV1 is cell autonomous, we studied mechano-sensing in isolated ASM cells at 32°C. Consistent with an earlier report (Mederos y Schnitzler *et al.*, 2008), we found that hypo-osmotic stretch rapidly increased intracellular [Ca^2+^] in ASM cells isolated from wild-type mice (Fig. 9A and B). In contrast, stretch evoked Ca^2+^ signals were significantly slower and smaller in ASM cells treated either with BCTC or isolated from TRPV1-null mice (Fig. 9A and B, peak rise *P* <0.001). Notably, reducing the temperature from 32°C to 23°C abolished stretch-evoked Ca^2+^ signaling in ASM cells (P < 0.0001, Fig. 9C).

**Figure 9.**
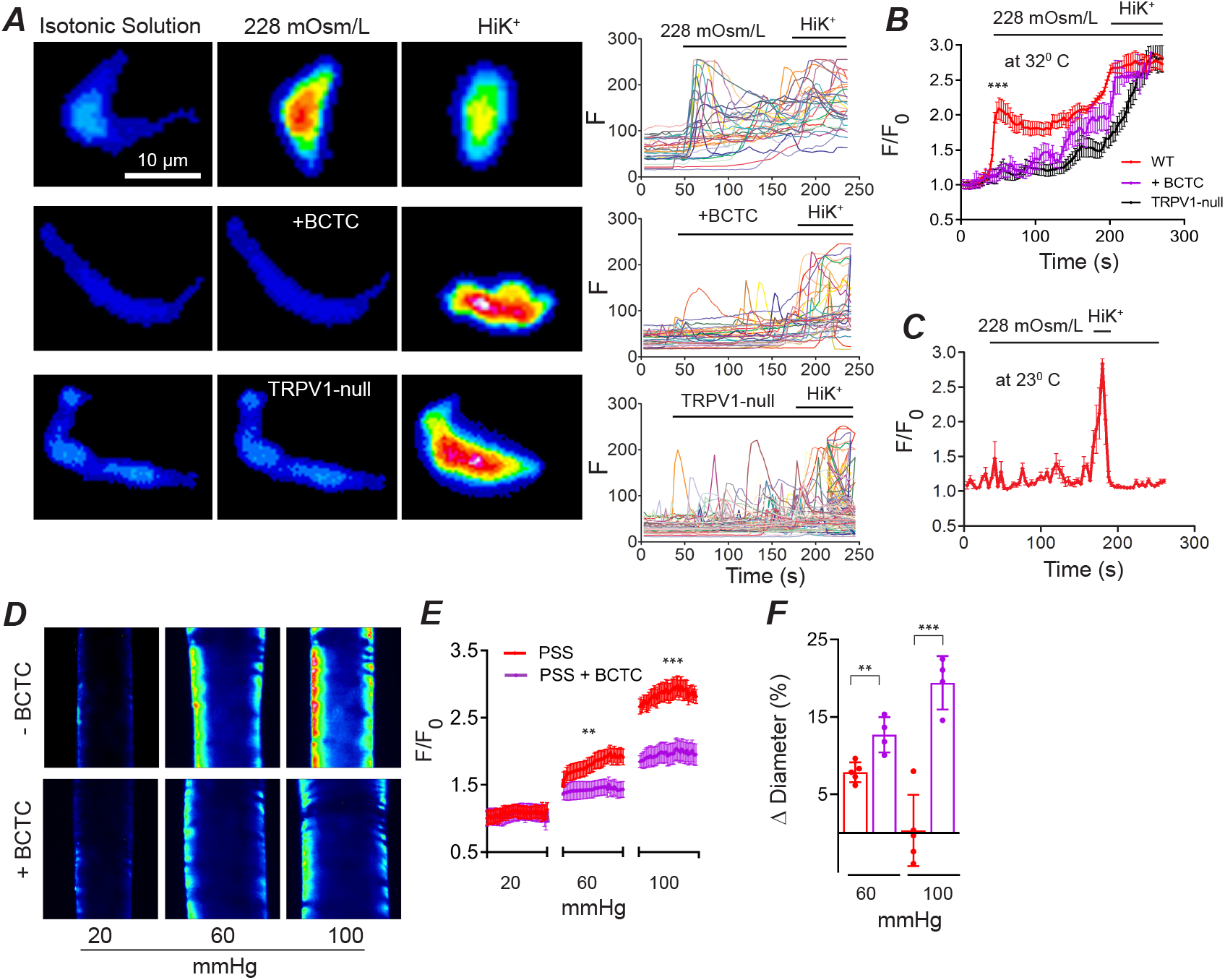
Stretch of arterial smooth muscle cells activates TRPV1 in a heat dependent manner. ***A-C***, Ca^2+^ signaling evoked by a hypo-osmotic solution in ASM cells isolated from wild-type (n = 47), wild-type plus BCTC (n = 35) and TRPV1-null (n = 30) mice. Both, BCTC (1 μM), and reducing temperature from 32°C to 23°C inhibits the stretch-evoked response (n = 28). ***D*** and ***E***, Intraluminal pressure evoked Ca^2+^ increases in arteries denuded of endothelium evoked are blocked by BCTC (3 μM; control, n = 51 and BCTC, n = 29). ***F*,** Changes in diameter of denuded arteries with or without BCTC (n = 4, unpaired t-test, \*\**P* = 0.0048, \*\*\**P* = 0.0002).

Osmotic stimuli are not perfect surrogates for mechanotransduction, therefore, to test the effects of physiological stretch we performed Ca^2+^ imaging in isolated pressurized arteries denuded of the endothelium. Under control conditions, increasing intraluminal pressure (from 20 to 60 and 100 mm Hg) triggered graded increases in Ca^2+^ accompanied by an unchanged vessel diameter (Fig. 9D and E). In contrast, BCTC inhibited the pressure-evoked rise in Ca^2+^ leading to vessel dilation (Fig. 9D and E). These results confirm that membrane stretch of vascular smooth muscle cells rapidly activates TRPV1.

Next, we explored the underlying mechanism for stretch-induced activation of TRPV1. Although not intrinsically mechanosensitive, TRPV1 is activated or sensitized by PLC-dependent signaling (Premkumar & Ahern, 2000; Chuang *et al.*, 2001; Tominaga *et al.*, 2001). Indeed, both PLCβ and PLCγ isoforms are implicated in membrane stretch and the generation of myogenic tone (Osol *et al.*, 1993; Matsumoto *et al.*, 1995; Gonzales *et al.*, 2014). We found that the PLC inhibitor, U73122, but not the inactive analogue decreased arterial tone to a similar extent as BCTC (Fig. 10). PLC cleaves PIP_2_ to form diacylglycerol (DAG). In turn, DAG can activate/sensitize TRPV1 channels via PKC-dependent phosphorylation (Premkumar & Ahern, 2000; Vellani *et al.*, 2001; Numazaki *et al.*, 2002). We found that the PKC antagonist, GX19203X, inhibited the development of tone in these arterioles to the same extent as inhibition of PLC. This result agrees with earlier studies demonstrating a critical role for PKC in the myogenic tone of skeletal muscle arterioles (Hill *et al.*, 1990; Korzick *et al.*, 2004; Hong *et al.*, 2016). Collectively, these results support a signaling pathway whereby membrane stretch engages a PLC/PKC pathway to activate TRPV1 (Fig. 10C). In turn, the resultant depolarization activates Ca_V_1.2 to trigger muscle contraction.

**Figure 10.**
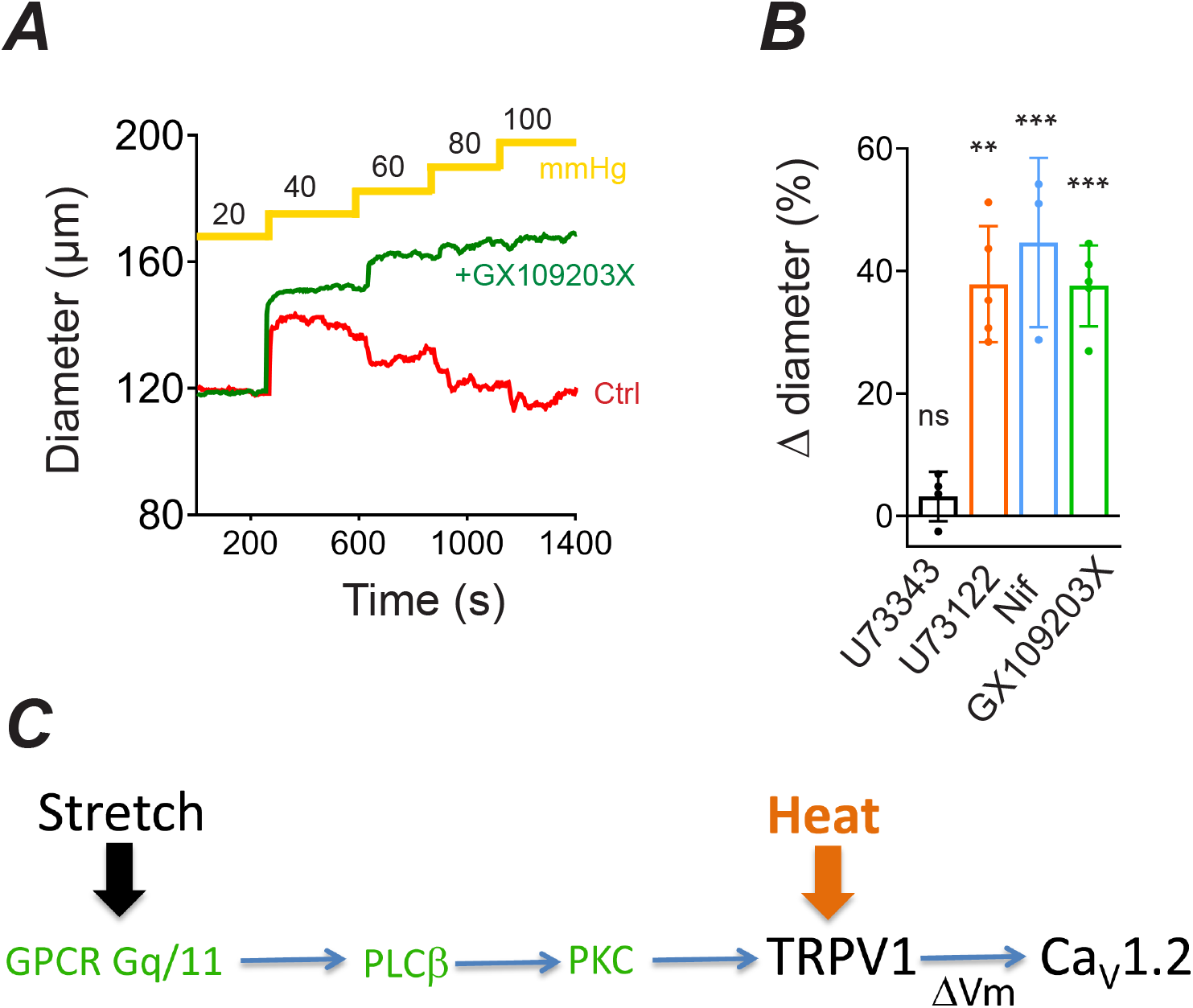
Arterial stretch activates TRPV1 in a PLC/PKC dependent manner. ***A***, Diameter versus pressure in an isolated skeletal muscle arteriole under control conditions (red) or with PKC inhibitor, GX109203X (10 μM, green). ***B***, Mean change in diameter of pressurized arteries (80 mm Hg, 34°C) treated with the PLC inhibitor, U73122 (10 μM, n = 4), the inactive analog, U73343 (10 μM, n = 4), nifedipine (n = 3) and GF 109203X (n=4), ***C***, Proposed signaling cascade for the generation of TRPV1-dependent myogenic tone.

## Discussion

Myogenic tone is a fundamental auto-regulatory property of arterioles that enables precise, local control of tissue perfusion. Although, fairly ubiquitous throughout the circulation there are differences in myogenic tone properties among tissues. Notably, the generation of myogenic tone is markedly faster in skeletal muscle (Hill *et al.*, 1990) and heart (Mosher *et al.*, 1964; Muller *et al.*, 1993) compared with the brain (Welsh *et al.*, 2002) and mesentery (Blodow *et al*., 2014). The underlying mechanisms for these differences are unclear. Our data show that TRPV1, localized in skeletal muscle, coronary and adipose arteries, participates in the regulation of myogenic tone. Several lines of evidence support this conclusion. Administration of the selective TRPV1 pharmacologic inhibitors, BCTC and AMG517, dilated arterioles *in vitro* and *in vivo* decreased blood pressure and increased coronary perfusion. The depressor effect of TRPV1 antagonists was transient due to an increase in HR. This cardiovascular compensation and/or inadequate dosage may explain why BP changes were not reported after oral administration of TRPV1 antagonists in humans and animals (Chizh *et al.*, 2007; Gavva *et al.*, 2007). Importantly, the antagonists failed to elicit responses in arteries from TRPV1-null mice confirming a selective action at TRPV1. Furthermore, BCTC inhibited the constriction of isolated arterioles in response to raised intraluminal pressure, and arterial smooth muscle cells in response to stretch. We found that myogenic tone in skeletal muscle arterioles (120 μm diameter) was largely sensitive to nifedipine (75-80%), indicating an important role for Ca_V_1.2 and in agreement with earlier findings (Knot & Nelson, 1998; Mauban *et al.*, 2013). BCTC blocked ~two-thirds of this nifedipine-sensitive tone, while the TRPM4 antagonist, 9-phenanthrol inhibited the remaining fraction. In contrast to pharmacologic blockade, genetic disruption of TRPV1 did not alter the steady-state tone measured 5 minutes after a pressure step. Developmental compensation by other ion channels in these constitutive TRPV1 knock-out mice may explain this discrepancy. Indeed, we found that expression of TRPM4 and TRPP1 increased significantly in arteries of TRPV1-null mice. Further, 9-phenanthrol elicited a greater inhibitory effect in TRPV1-null arteries consistent with the larger fraction of TRPM4-dependent tone. Although these data should be interpreted cautiously; 9-phenanthrol may not be perfectly selective for TRPM4 (Burris *et al.*, 2015), the results support the hypothesis that TRPM4 mediates the TRPV1-independent tone in WT arteries, and that upregulation of TRPM4 expression partly compensates for genetic disruption of TRPV1. Potentially other TRP channels, including TRPP1, may also contribute to restore steady-state tone in TRPV1-null mice.

A key finding of this study is that TRPV1 regulates the rate of myogenic tone. Although TRPV1-deficient arteries exhibited unchanged steady-state tone, the development of tone was slowed ~2-3-fold compared with wild-type arteries. This difference was most pronounced in the smallest diameter arterioles studied (25 μm diameter) where disruption of TRPV1 expression increased the T_1/2_ for development of tone from ~15 to 45 s. The rapid tone in WT arteries agrees with earlier reports. For example, raised intraluminal pressure in third-order cremaster muscle arterioles arterioles (~10-15 μm diameter) triggered a decrease in diameter within 10 s (Hill *et al.*, 1990). Similarly, rapid myogenic responses are evident in isolated coronary arterioles (Muller *et al.*, 1993) and in the beating dog heart (Mosher *et al.*, 1964), where increases in perfusion pressure triggered autoregulation of coronary flow within 10 s.. Although we can’t exclude contribution of other factors, our data suggest that the abundant expression of TRPV1 in skeletal muscle and coronary arterioles significantly contributes to the rapid myogenic tone of these tissues. In addition to controlling the onset of myogenic tone, our data reveal a critical role for TRPV1 in the rapid cessation of tone. Skeletal muscle arterioles characteristically dilate at the beginning of muscle contractions (Clifford, 2007). Indeed, using intravital imaging we observed prominent vasodilation following arteriole constrictions of duration from 4 to 40 s. Notably, BCTC completely occluded this reactive vasodilation. Furthermore, genetic disruption of TRPV1 inhibited greater than 70% of the response demonstrating minimal developmental compensation. Metabolic factors may underlie reactive hyperemia particularly with long duration vessel occlusion. However, previous studies have shown that even very brief (100 ms) muscle contractions elicit vasodilation of skeletal muscle arterioles *in situ* (Sinkler *et al.*, 2016). Further, rapid onset vasodilation occurs in isolated arteries (Clifford *et al.*, 2006) and in the human forearm (Kirby *et al.*, 2007) in response to intermittent raised extravascular pressure. Thus, rapid onset vasodilation may purely reflect an effect of mechanical compression of the vessel. Importantly, our findings suggest that deactivation or inactivation of TRPV1, that disrupts myogenic tone, is a critical component of this process.

How does TRPV1, not recognized as intrinsically mechanosensitive, participate in arterial mechano-signaling? Gq/G11 GPCRs are candidate cellular mechanosensors and are implicated in the detection of intraluminal stretch (Mederos y Schnitzler *et al.*, 2008). In skeletal muscle arterioles, the type 1a angiotensin receptor (AT_1a_R) contributes to the generation of myogenic tone, signaling via a PLC-PKC pathway (Hong *et al.*, 2016). Indeed, we found a key role for PLC/PKC signaling; pharmacological antagonists of these proteins inhibited myogenic tone by 75-80%. Notably, this same fraction of myogenic tone was sensitive to nifedipine. Thus, consistent with earlier reports (Hill *et al.*, 1990; Korzick *et al.*, 2004), PKC signaling underlies the majority of the myogenic tone of skeletal muscle arterioles, linking membrane stretch to the activation of voltage–gated Ca_V_1.2 channels. Our data suggest that TRPV1 is a key intermediary in this signaling. Notably, PKC-dependent phosphorylation sensitizes TRPV1 to various stimuli including heat, reducing the temperature threshold substantially from 42°C to 32°C (Sugiura *et al.*, 2002). Further, the diverse chemical and physical sensor domains in TRPV1 are allosterically coupled to channel gating (Tominaga *et al.*, 1998; Brauchi *et al.*, 2004; Voets *et al.*, 2004; Matta & Ahern, 2007). Thus, PKC in combination with other stimuli may underlie rapid activation of TRPV1 and changes in myogenic tone. Conversely, a fall in intraluminal pressure and de-phosphorylation of TRPV1 would rapidly decrease TRPV1 activity and reduce tone. Interestingly, PKC also stimulates TRPM4 (Guinamard *et al.*, 2002; Nilius *et al.*, 2005) that may be responsible for the non-TRPV1 fraction of myogenic tone inhibited by 9-phenanthrol. Thus, PKC stimulation of TRPV1, and to a lesser extent TRPM4, mediates depolarization-induced activation of Ca_V_1.2. Previously, we showed that Ca^2+^ permeability through TRPV1 activated by high concentrations of capsaicin is sufficient to constrict arterioles (Phan *et al.*, 2020). Here we found that nifedipine completely inhibits myogenic tone, negating a contribution of Ca^2+^ entry through TRPV1. This may reflect weak activation of TRPV1 during stretch signaling compared with capsaicin. Moreover, the relative level of TRPV1 expression in vascular smooth muscle is low, ~10% of sensory neurons (Phan *et al.*, 2020), and therefore a high open probability of these channels is likely required to sufficiently raise myoplasmic Ca^2+^.

A prevailing hypothesis is that myogenic autoregulation holds arteries in a partly constricted state thereby facilitating bi-directional changes in vessel caliber and blood flow. Indeed, its suppression with ageing (Lott *et al.*, 2004; Ghosh *et al.*, 2015) and in diabetes/cardiovascular disease (Petersen *et al.*, 2002; Hodnett & Hester, 2007; Duncker *et al.*, 2015) underscores an important role for the myogenic response in cardiac and exercise performance. Our data reveal that TRPV1 speeds both the development and removal of myogenic tone and thus aids in the dynamic regulation of perfusion in skeletal muscle and the heart. This property may be advantageous in these tissues that experience both large and fast fluctuations in blood pressure. Further, our data reveal a critical role for TRPV1 in rapid reactive vasodilation, providing a molecular mechanism for how skeletal muscle contractions at the onset of activity immediately increase tissue blood flow. We propose that TRPV1 may enable a similar process in myocardial perfusion as our data show that TRPV1 contributes significantly to coronary myogenic tone.

## Author contributions

T.P designed and performed most of the experiments, analyzed the data and helped write the manuscript; G.A. conceived and designed experiments, performed electrophysiology and wrote the manuscript; H.T. performed experiments; N.S. assisted with mouse BP measurements and helped write the manuscript; H.G and R.B. performed rat physiology and helped write the manuscript. A.T. helped design experiments and helped write the manuscript. R.R. and M.K. performed coronary flow experiments.

